# How the initiating ribosome copes with (p)ppGpp to translate mRNAs

**DOI:** 10.1101/545970

**Authors:** Daria S. Vinogradova, Pavel Kasatsky, Elena Maksimova, Victor Zegarra, Alena Paleskava, Andrey L. Konevega, Pohl Milón

## Abstract

During host colonization, bacteria use the alarmone (p)ppGpp to reshape its proteome by acting pleiotropically on RNA and protein synthesis. Here, we elucidate how the translation Initiation Factor 2 (IF2) senses the cellular ppGpp to GTP ratio and regulates the progression towards protein synthesis. Our results show that the affinity of GTP and the inhibitory concentration of ppGpp for 30S-bound IF2 vary depending on the programmed mRNA. Highly translated mRNAs enhanced GTP affinity for 30S complexes, resulting in fast transitions to elongation of protein synthesis. Less demanded mRNAs allowed ppGpp to compete with GTP for IF2, stalling 30S complexes until exchange of the mRNA enhances the affinity for GTP. Altogether, our data unveil a novel regulatory mechanism at the onset of protein synthesis that tolerates physiological concentrations of ppGpp, and that bacteria can exploit to modulate its proteome as a function of the nutritional shift happening during infection.

## Main Text

During colonization, pathogenic bacteria reshapes its transcriptome and proteome to activate virulence genes, promote tissue-associated biofilm and enter dormancy, ultimately increasing aggressiveness, antibiotic tolerance, and persistence of the pathogen (reviewed in^1,2^). Host colonization entails fluctuations of nutrient availability that, generally, triggers stringent response in bacteria (Fig. 1). Stringent response is mediated by the ribosome-associated RelA/SpoT homolog protein superfamily and triggers the accumulation of the hyperphosphorylated guanosine nucleotides, altogether called (p)ppGpp (Fig. 1)^3^. Impaired (p)ppGpp production results in antibiotic sensitivity, low biofilm formation, and low pathogenicity of several bacteria, making the stringent response an appealing target for antibiotic development^4–6^. mRNA translation to proteins requires the action of several guanosine nucleotide-binding factors, on and off the ribosome. Although (p)ppGpp have been shown to bind translational GTPases^7,8^, the extent of inhibition and subsequent effect on protein synthesis regulation remain controversial.

**Fig. 1.**
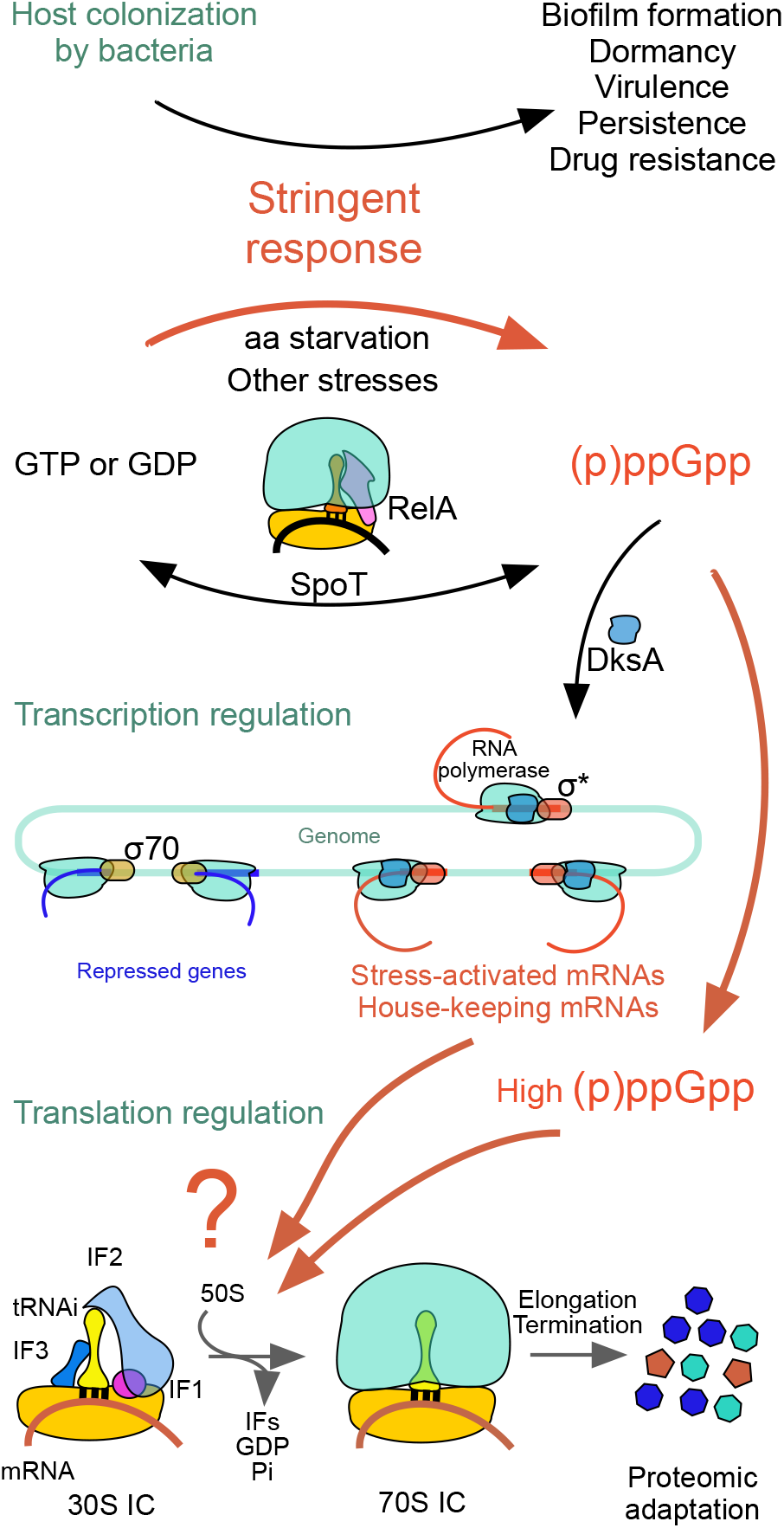
Bacterial stringent response and (p)ppGpp-mediated inhibition of translation initiation.

The initiating ribosome orchestrates a complex equilibrium between three initiation factors (IFs), mRNA and initiator tRNA (fMet-tRNA^fMet^, from here in tRNAi) to maximize the speed and accuracy of start codon selection, ultimately defining the reading frame for mRNA translation^9^. The translational GTP binding factor IF2 recruits tRNAi to the 30S pre-initiation complex (pre-IC), accompanies the subsequent isomerization to the 30S IC (start codon recognition), promotes the association of the large 50S subunit and occupies all intermediates of the 70S pre-IC leading to a ready-to-elongate 70S IC^9^. Although GTP has been shown to enhance IF2 activity, hydrolysis of GTP appears to be dispensable^10^. IF2 shows a broad spectrum of binding properties as a function of the bound guanosine nucleotide, enabling cycling between high and low affinity states: GTP to assemble the 30S IC, whereas GDP allows dissociation of the factor from the 70S pre-IC^11,12^. Both, the dispensability of GTP hydrolysis and the wide range of affinities displayed by the factor, allowed to propose IF2 to function as a molecular sensor of the stringent response. Indeed, ppGpp was shown to bind IF2, and to inhibit translation initiation and first peptide bond formation^8^. Whether (p)ppGpp stringently or permissively arrests translation initiation remained an open question. (p)ppGpp-dependent activation of genes and involvement in controlling the growth rate of the cell by acting in DNA, RNA and protein synthesis^13–16^argues for a permissive mechanism. Here, we use advanced fluorescence spectroscopy techniques to investigate how the initiating ribosome copes with (p)ppGpp accumulation to translate mRNAs, allowing cell survival and reshaping the bacterial proteome for environmental adaptation.

## Results

### Guanosine nucleotides as co-factors of IF2

We used Microscale Thermophoresis (MST) to measure the binding of fluorescently labelled initiator fMet-tRNA^fMet^ (Bpy-tRNAi) to 30S subunits as an indicator of 30S IC formation. MST allows to measure fluorescence changes derived from molecular drifts resulting from small equilibrium perturbations^17^ Upon equilibrium perturbations, the migration patterns of Bpy-tRNAi are related to bound and unbound states, allowing to determine dissociation constants of the interaction (see supplementary information: Fig. 1, 2, and experimental approach)^17,18^. Formation of 30S ICs resulted in an increase of thermophoresis as compared to the free Bpy-tRNAi. On the contrary, a decrease in thermophoresis indicated the dissociation of Bpy-tRNAi from 30S complexes.

**Fig. 2.**
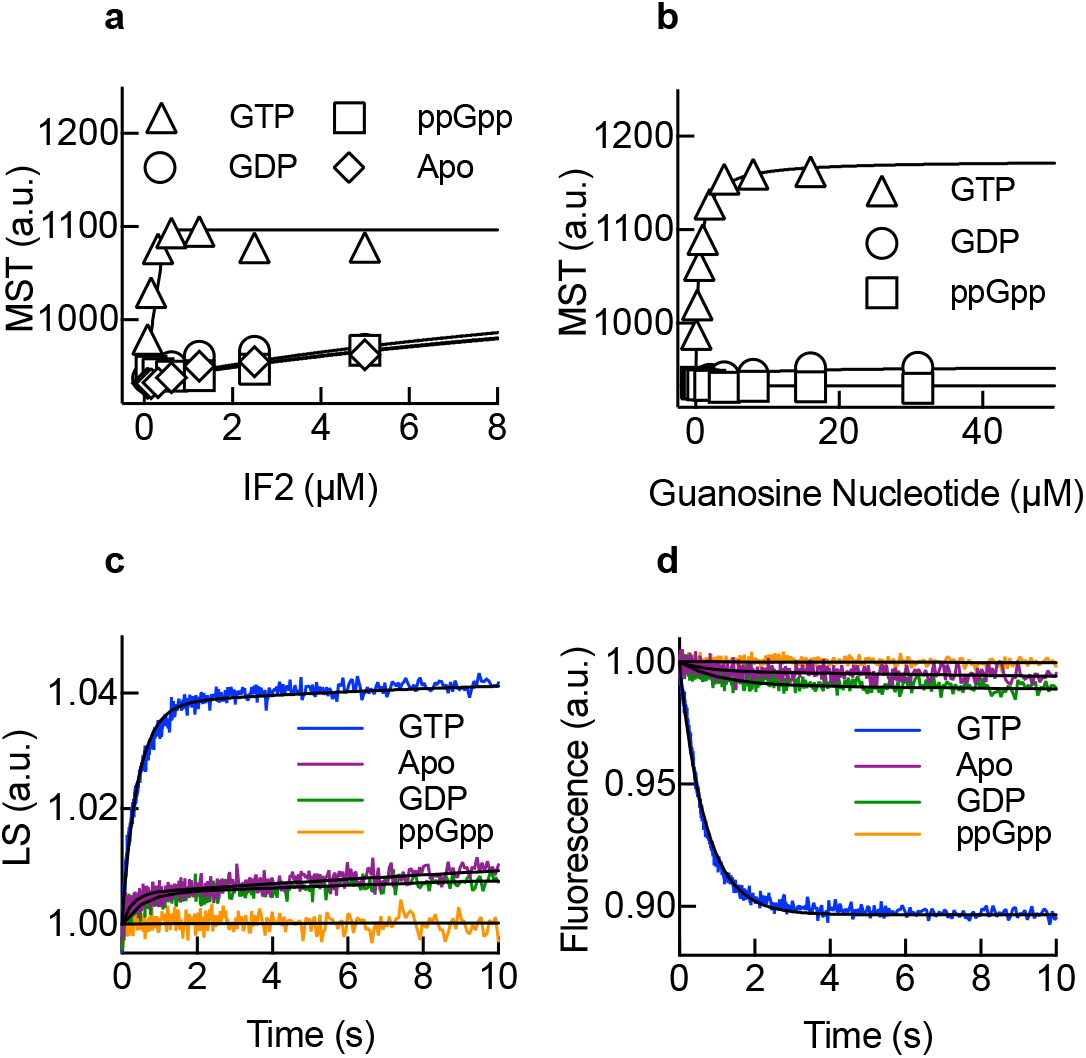
ppGpp-mediated inhibition of translation initiation. **(a)** 30S IC formation measured by MST at increasing concentrations of IF2 as a function of ppGpp (squares), GTP (triangles), GDP (circles) or in the absence (diamonds) of a guanosine nucleotide. Continuous lines show best fits using a Hill equation. **(b)** as (a) to measure the concentration dependence for each of the tested guanosine nucleoside. Continuous lines indicate non-linear regression fittings. 3-5 measurements were performed, mean and standard deviation are plotted. **(c)** 70S pre-IC formation as measured by light scattering (LS) on a stopped-flow apparatus. 30S initiation complexes were formed in the absence of (purple) or the presence of 0.5 mM ppGpp (orange), GTP (blue) or GDP (green) and rapidly mixed with 50S subunits^20^. Continuous lines show best fits using an exponential function for two steps. **(d)** 70S IC formation as measured by Bpy-tRNAi accommodation in the P site of the ribosome^12^. 30S ICs were formed with Bpy-tRNAi and rapidly mixed with 50S subunits on a stopped-flow apparatus. Colours are as in (c). Continuous lines show best fits using an exponential function for a single reaction step. All reactions are mean values of five to ten replicates.

First, we measured 30S IC formation as a function of IF2 concentration, in the presence of different guanosine nucleotides. Addition of GTP increased the amplitude of thermophoresis stoichiometrically with IF2, while in the presence of GDP, ppGpp or without any nucleotide (Apo), the amplitude of thermophoresis was lower (Fig. 2a), in agreement with IF2 requiring GTP for rapid recruitment of tRNAi to the 30S complex^19^. EC_50_ binding concentrations for Bpy-tRNAi were very low for IF2 bound to ppGpp, GDP or in the absence of any guanosine nucleotide (Apo) and coincided with measurements of IF2-Bpy-tRNAi ternary complex formation in the absence of the 30S subunit (Supplementary Fig. 3). Thus, ppGpp and GDP may program IF2 to de-stabilizes tRNAi on the 30S complex, precluding 30S IC formation. Consistently, guanosine nucleotide titrations showed thermophoresis increase for GTP and to a lesser extent for GDP, whereas ppGpp failed to promote Bpy-tRNAi binding at any concentration (Fig 2b). GTP seems to activate IF2 to promote tRNAi binding to the 30S IC, whereas, GDP and ppGpp prevent IF2 from recruiting the initiator tRNA.

To investigate the transition to translation elongation as a function of IF2-bound nucleotides, we monitored the formation of the 70S pre-IC by scattered light (LS) and 70S IC formation by Bpy-tRNAi accommodation by fluorescence using stopped-flow techniques^12,21^(Fig. 2c,d). Rapid mixing of 50S subunits with 30S complexes formed with GTP resulted in a rapid increase of LS over time (*k*_app_ = 2.9 ± 0.3 s^−1^), indicating that 70S pre-ICs are readily formed (Fig. 2c). Bpy-tRNAi showed a rapid decrease in fluorescence over time following 50S subunit joining (*k*_app_ = 1.7 ± 0.2 s^-1^) (Fig. 2d). Altogether, GTP programs IF2 to form a 30S IC that is capable of rapid binding the 50S subunit and accommodating tRNAi in the 70S IC to accept the incoming elongator aminoacyl tRNAs. Interestingly, the omission of nucleotides or presence of GDP showed similar rates for both reactions, albeit with much lower efficiencies (amplitude) than that observed with GTP. ppGpp, instead, drastically reduced the rates and extent of 50S subunit joining and Bpy-tRNAi accommodation, an indicator that ppGpp acts at a prior step of the translation initiation pathway, i.e., before 30S IC formation. Although both, GDP and ppGpp, could compete with GTP for IF2, cellular concentrations of GDP are low at any cell growth condition^22^, whereas (p)ppGpp accumulate to millimolar ranges during bacterial stringent response^22,23^. Overall, guanosine nucleotides act as co-factors of IF2, modulating its capacity to position the initiator tRNA along the pathway of translation initiation (GTP vs. GDP) or as a sensor of the stringent response (GTP vs. ppGpp).

### mRNA dependence of ppGpp inhibition

mRNAs may contain the determinants to enter translation at otherwise inhibiting concentrations of ppGpp, allowing GTP to compete with ppGpp (Fig. 3a). To test this model, we used two house-keeping mRNAs, coding for the essential proteins EF-Tu (*mTuf*A) and IF1 (*mlnf*A), with the former being 40-fold more expressed than the latter in *E. coli*^24^. 30S IC formation as a function of mRNA concentrations was higher for *mTuf*A than for *mlnf*A or the model mRNA mMF1 as evaluated from the thermophoresis amplitude (Supplementary Fig. 4 and Tables 1-3). Similarly, GTP titrations in complexes formed with each mRNA showed the same thermophoresis trend for 30S IC formation: mTu/A>mInfA>mMF1 (Fig. 3b). Additionally, GTP dissociation constants differed around an order of magnitude as a function of the mRNA used. The calculated *K_D_* for the 30S IC programmed with *mTuf*A was about 7-fold lower (*K*_D_ = 0.28 ± 0.04 μM) than that for *mlnf*A (*K*_D_ = 1.9 ± 0.2 μM) and about 2-fold lower than that for mMF1 (*K*_D_ = 0.65 ± 0.05 μM) (Fig. 3c). Thus, mTufA, which codes for the highly abundant EF-Tu, possesses functional determinants that allow efficient 30S IC formation with IF2 strongly binding GTP. On the other hand, *mlnf*A results in less efficient 30S IC formation and weaker binding of GTP.

**Fig. 3.**
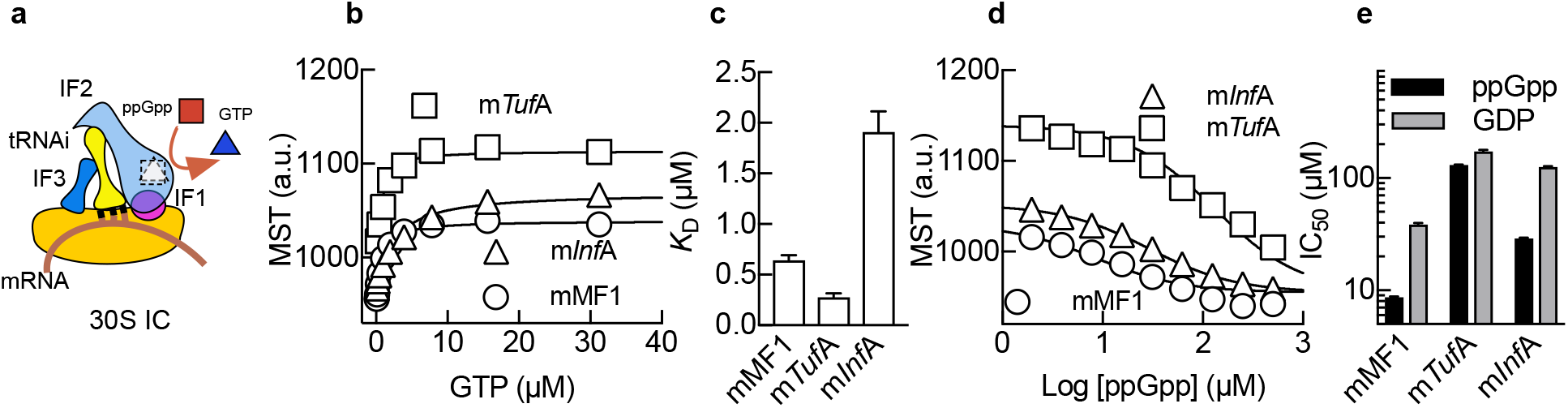
mRNA dependence and MST analysis of 30S IC formation. **(a)** Scheme of the 30S IC highlighting ppGpp (red square) and GTP (blue triangle) competition for IF2. **(b)** 30S IC formation with increasing concentrations of GTP for *mTuf*A (squares), *mlnf*A (triangles) or model messenger mMF1 (circles). 30S IC formation was measured by MST and analysed as described above (Fig. 2). **(c)** Comparison of *K*_D_ calculated from (b). **(d)** ppGpp to GTP competitive assays for 30S IC formation. 30S complexes formed with each mRNA (symbols as in b) and in the presence of 50 μM GTP were subjected to increasing concentrations of ppGpp. Log of competitor concentrations is plotted and used for determining the inhibitory concentration for 50% inhibition (IC_50_) using a same-site competition model. **(e)** Bar graph comparing IC_50_ values for ppGpp (black) or GDP (grey) for all three mRNAs. Continuous lines indicate nonlinear regression fittings. 3-5 measurements were performed, mean and standard deviation are plotted.

ppGpp competition experiments with GTP showed a decrease in 30S IC formation for all mRNAs with increasing concentrations of ppGpp, however with different dependencies on the competing nucleotide (Fig. 3d). The calculated inhibitory concentrations (IC_50_) ranged over 15-fold with *mTuf*A being the least sensible to ppGpp (IC_50_ = 132 ± 1 μM), mMF1 the most sensible (IC_50_ = 8.6 ± 0.2 μM), and *mlnf*A being inhibited with an intermediate concentration (IC_50_ = 29 ± 1 μM) (Fig. 3e). Similar experiments performed with GDP required higher concentrations of the competitor to inhibit 30S IC; however, mRNAs dependencies were maintained (Fig. 3e, supplementary Fig. 5). Thus, ppGpp-mediated inhibition of translation initiation is dynamic and dependent on the mRNA bound to the 30S complex, with the highly translated *mTuf*A being more tolerant of ppGpp concentrations than *mlnf*A.

### ppGpp halts 70S IC formation

Late events of translation initiation entail the association of 30S ICs with the large ribosomal subunit 50S leading to the intermediate 70S pre-IC, and after IFs dissociation results in a ready-to-elongate 70S IC (Fig. 4a)^12^. Here, we measured the velocities of 70S pre-IC formation as a function of the bound mRNA, the IF2-bound nucleotide, and the ability of ppGpp to compete with GTP (Fig. 4). Rapid kinetics analysis of the 50S joining to 30S ICs show that complexes programmed with *mTuf*A resulted in higher 70S pre-IC formation than those formed with *mlnf*A or mMF1, essentially following the same trend as observed by thermophoresis analysis (Fig. 4b). Addition of ppGpp as a competitor for GTP showed an overall decrease of 70S pre-IC formation efficiency for all mRNAs; however, complexes harbouring *mTuf*A appeared least affected while *mlnf*A were the most affected (Fig. 4c). Using GDP as a competitor for GTP also resulted in a decreased efficiency of 70S pre-IC formation; however, inhibiting the reaction to a lesser extent than ppGpp (Fig. 4d, supplementary Fig. 6). Additionally, the formation of the 70S pre-IC appears to be kinetically influenced by the *mlnf*A showing 3 to 4-fold slower velocities than *mTuf*A or mMF1 (Fig. 4e). However, the nucleotide competing with GTP, either GDP or ppGpp, did not perturb the initial rate of the 50S joining to 30S ICs, suggesting that the observed fraction corresponds to 30S ICs containing GTP (Fig. 4e). Thus, ppGpp competition with GTP results in fewer 30S ICs habilitated to recruit the 50S subunit; albeit, mRNAs modulate the GTP-bound fraction. Similar reactions in the absence of GTP resulted in negligible rapid 70S pre-IC formation for either ppGpp and some transitions for GDP or without any nucleotide (Supplementary Fig. 7).

**Fig 4.**
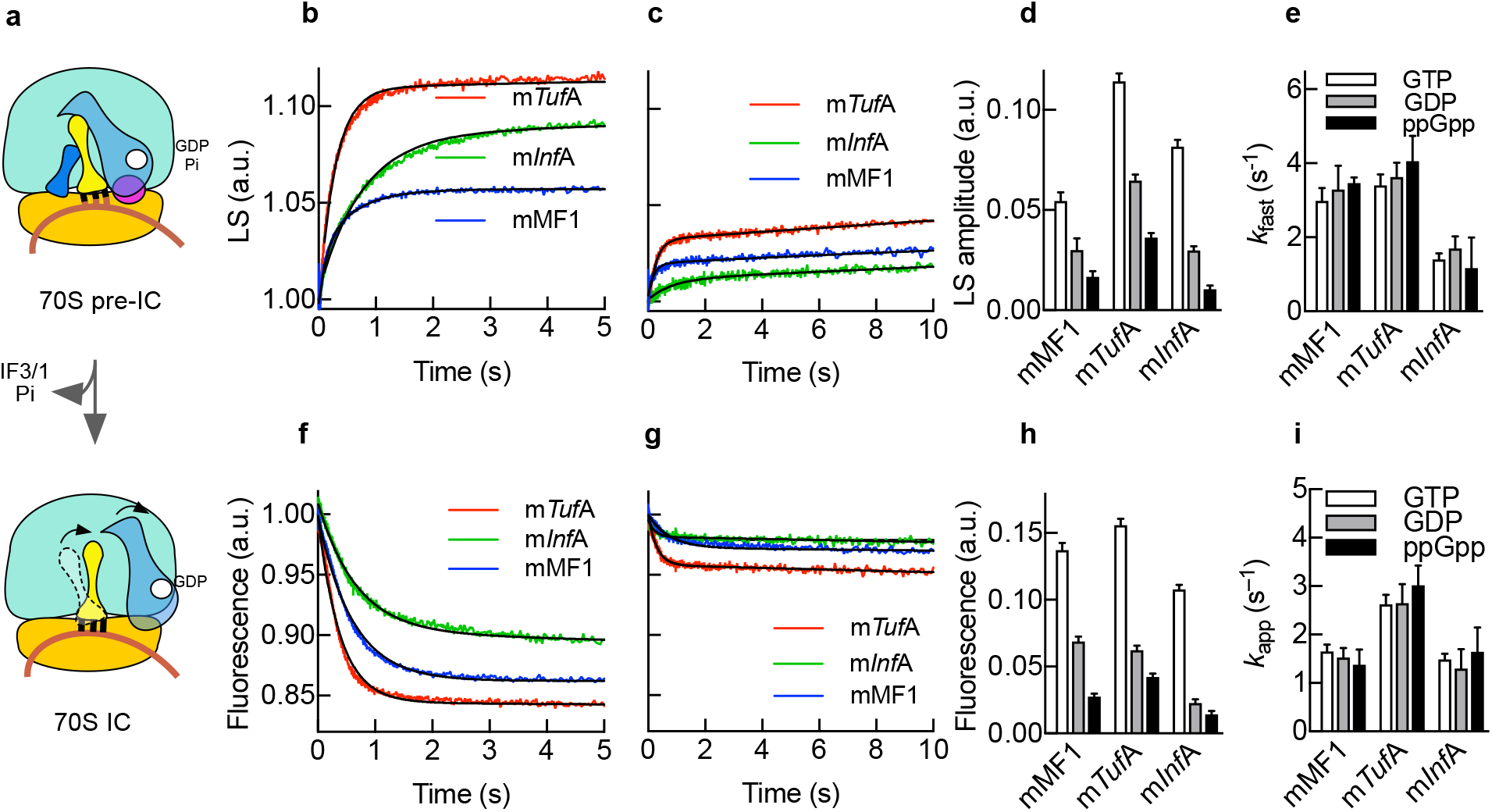
Kinetic parameters of ppGpp-mediated regulation of 70S IC progression. **(a)** Scheme of 70S IC formation. 30S complexes programmed with *mTuf*A (red), *mlnf*A (green) or mMF1 (blue) and 20 μM GTP and the absence **(b)** or presence **(c)** of 200 μM ppGpp were reacted with 50S subunits. Time traces were analysed by non-linear regression with two exponential terms. **(d)** Bar graph comparing amplitudes in the absence of any competing nucleotide (white) or in the presence of ppGpp (black) or GDP (grey). **(e)** Bar graph comparing apparent rates of 70S pre-IC formation (colours as in d). 70S IC formation as measured by Bpy-tRNAi accommodation in the absence **(f)** or presence of 200 μM ppGpp **(g)**. Time traces were analysed by non-linear regression with one exponential term. **(h)** Bar graph comparing amplitude variations as a function of mRNAs and guanosine nucleotides (colours as in d). **(i)** Bar graph comparing apparent rates of 70S IC formation for all three mRNAs (bar colours as in d). Continuous lines show best fits using an exponential function for a single reaction step. All time traces are mean values of five to ten replicates, mean and standard deviation are plotted.

IF2 populates every intermediate of the multi-step reaction leading to 70S IC formation, ultimately promoting the accommodation tRNAi in the P site before factor dissociation (Fig. 4a)^12,25^. 70S IC formation was assessed by measuring Bpy-tRNAi accommodation using the stopped-flow technique as a function of the guanosine nucleotide competing with GTP for all three mRNAs. In the absence of any GTP competitor, tRNAi accommodated rapidly to the complexes after the 50S joining (Fig. 4f), with overall efficiencies reflecting those of 30S IC formation as measured by thermophoresis (Fig. 3b) or 70S pre-IC as measured by light scattering (Fig. 4b). Replacement of GTP by GDP or ppGpp resulted in small fluorescence changes for tRNA accommodation, indicating very few 30S complexes contained the tRNAi (Supplementary Fig. 8). Addition of GDP or ppGpp as competitors of GTP resulted in a defined decrease of overall efficiencies of tRNAi accommodation; however, ppGpp showed a higher degree of inhibition than GDP (Fig. 4g, supplementary Fig. 6). As observed for 70S pre-IC formation, complexes programmed with *mTuf*A appeared to be less sensitive to ppGpp while those programmed with *mlnf*A showed more susceptibility (Fig. 4h). Non-linear analysis of the time dependencies shows that the tRNAi accommodates at different rates for each mRNA. Although the extent of the reaction is affected by the competing nucleotide, the apparent rates of tRNAi accommodation appeared not to be influenced, indicating that the resulting amplitude corresponds to GTP-bound 30S ICs (Fig. 4i). Altogether, the formation of 70S ICs are halted by ppGpp. The progression towards protein synthesis elongation is mediated by the competition between GTP with ppGpp during 30S IC formation in an mRNA-dependent manner.

### Permissive mRNA translation at high ppGpp

Our results are consistent with IF2 sensing ppGpp to GTP ratios in an mRNA-dependent manner and ultimately translating them into protein output efficiencies (Fig. 5a). To test this premise, we used a cell-free translation system at physiological concentrations of ribosomes, aminoacyl-tRNAs, translational GTPases, GTP and varying ppGpp concentrations (up to 4 mM)(Fig. 5). In the absence of ppGpp, translation of *mTuf*A was three-fold higher than *mlnf*A (Fig. 5b), consistent with our results obtained by measuring every previous step, 30S IC, 70S pre- and IC (Fig. 3,4). Addition of ppGpp resulted in decreased translation efficiencies for both mRNAs in a ppGpp concentration-dependent manner (Fig. 5c). The inhibitory concentration IC_50_ differed for each mRNA, with *mTuf*A being 4-fold more tolerant to ppGpp than *mlnf*A (Fig. 5d).

**Fig. 5.**
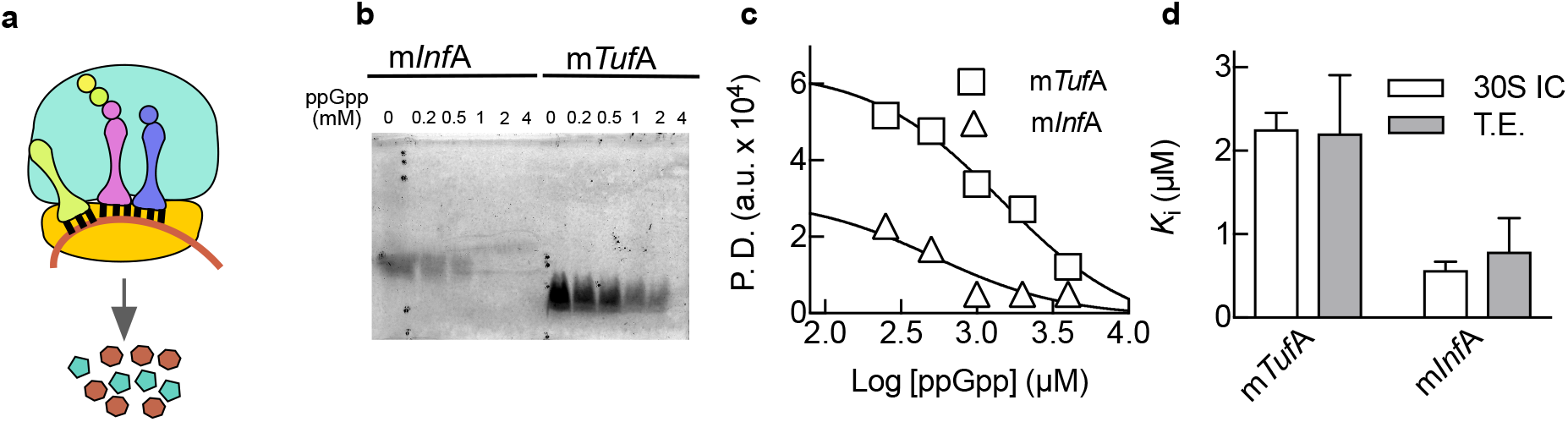
ppGpp-mediated inhibition of mRNA translation. **(a)** Scheme of the translating 70S complex. **(b)** *In vitro* translation of *mTuf*A and *mlnf*A derivatives harbouring a coding sequence for the Lumio labelling system (see Methods). Protein synthesis reactions were started in the presence of 2 mM GTP and increasing concentrations of ppGpp. Synthetized proteins were fluorescently labelled and resolved by 20% SDS-PAGE. The resulting images were analysed by pixel densitometry using ImageJ^26^ to estimate efficiencies of translation (T.E.). (c) Log of ppGpp concentrations was plotted and used for determining the inhibitory concentration for 50% inhibition (IC50) using a same-site competition model. (d) Comparison of inhibitory constants for both mRNAs as measured during 30S IC formation (white) or overall translation efficiency (grey). Mean and standard deviation from 3-5 measurements are plotted.

The calculated affinity of GTP (Fig. 2) for complexes differing on the programmed mRNA allowed to estimate the ppGpp inhibitory constants for each corresponding 30S IC and translation efficiencies. The calculated ppGpp inhibitory constants for translation efficiency were similar to those obtained for 30S IC formation as measured by thermophoresis, reaffirming IF2 as the primary target for ppGpp-mediated regulation during protein synthesis (Fig. 5d)^8^. Remarkably, *mTuf*A was translated at ppGpp concentrations that have been reported during cell starvation^22^, indicating that the protein synthesis apparatus is capable of tolerating high concentrations of the alarmone. Altogether, the protein synthesis apparatus can translate mRNAs at physiological concentrations of ppGpp; however, the initiating ribosome is able to sort which mRNAs shall enter the elongation phase of protein synthesis.

## Discussion

Our results provide unprecedented details on the dynamic regulation of the initiating ribosome and allowed us to untangle a novel mechanism that allow bacteria to cope with mRNA translation during stringent response. The canonical model suggests that upon (p)ppGpp accumulation the translation machinery halts until more favourable growth conditions are available. Shutdown of protein synthesis during stringent response was supported by several reports indicating (p)ppGpp binds and inhibits translational GTPases^7,27^. However, the canonical model fails to explain how a subset of proteins are synthetized during overexpression of RelA, the primary (p)ppGpp synthetizing factor^28^. Additionally, (p)ppGpp activates the transcription of a number of genes. How these mRNAs are translated remained unexplained. More recent reports showed that (p)ppGpp are not restricted to stringent response, but their concentration fluctuates as a function of growth rate^13^. A tight RNA/protein and DNA/protein synthesis coordination in *E. coli* was shown to be regulated by (p)ppGpp^13^. On the other hand, the rate of protein elongation appears to be unaffected during stationary phase (starvation), characterized by high levels of the alarmone^29^. Thus, (p)ppGpp-mediated inhibition of protein synthesis appears to be permissive rather than a strict on/off mechanism. Our results support a permissive mechanism where IF2 translates the nutritional availability of the bacterial cell into protein synthesis efficiencies by sensing the ppGpp to GTP ratios (Fig. 6).

**Fig. 6.**
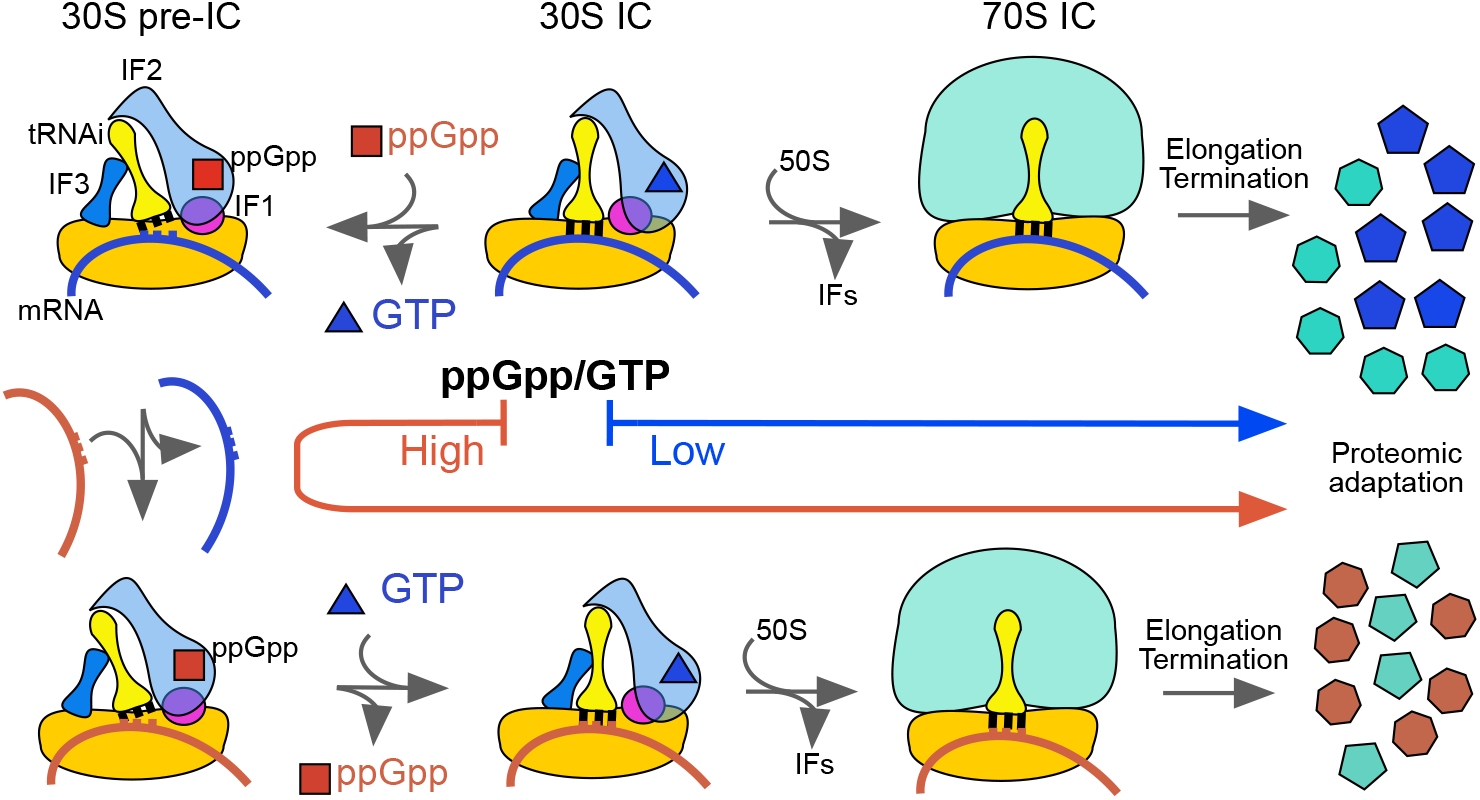
30S-bound IF2 senses the stoichiometry of ppGpp to GTP. At low ppGpp/GTP ratios IF2 binds GTP and can rapidly transit towards translation elongation (blue arrow). At high ppGpp/GTP ratios ppGpp binds IF2, precluding start codon recognition and promoting the 30S pre-IC. In turn, the 30S pre-IC can exchange the bound mRNA for a more ppGpp-tolerable transcript, allowing GTP to replace the tetraphosphate and proceed to protein elongation.

ppGpp accumulation results in IF2 modulating the 30S pre-IC to IC equilibrium, ultimately contributing to define the efficiency by which the mRNA enters the elongation phase of protein synthesis (Fig 6). ppGpp-bound IF2 shifts the translation initiation equilibrium towards the liable 30S pre-IC in an mRNA dependent manner. The potential of each mRNA to be translated arises from the intrinsic capability of mRNAs to program 30S ICs (Fig. 3a, supplementary Fig. 4), the GTP affinity for IF2-bound complexes (Fig. 3c), and the tolerance to ppGpp (Fig. 3e). We observe that the highly translated *mTuf*A mRNA is more tolerant to ppGpp than *mlnf*A despite that both code for house-keeping and essential proteins. *mTuf*A initiates translation 2 to 4-fold more efficiently than *mlnf*A in the absence of any competing mRNA. In a cellular context, the observed difference may be accentuated to the extent observed *in vivo* (40-fold,^24^) due to the availability of free 30S subunits and competing mRNAs. On the other hand, the GTP affinity for 30S ICs programmed with *mTuf*A is 7-fold higher if compared to *mInfA,* both in the low micromolar range. In contrasts, previous studies reported two to three orders of magnitude lower affinities between free IF2 and GTP^7^ Thus, the formation of the 30S IC entails an affinity gain for GTP to the 30S-bound IF2 of at least two orders of magnitude. This increase of affinity is of particular importance in the context of stringent response where GTP concentration have been shown to drop to micromolar ranges^22^. However, the affinity gain of the 30S-bound IF2 for GTP does not prevent ppGpp to compete. Altogether, IF2 may act as a molecular sensor for cellular homeostasis, coupling functional determinants of the mRNA to ppGpp concentration to cope for the environmental adaptation of bacteria during infection.

## Methods

### Biological preparations

#### Ribosomalsubunits

30S subunits were prepared from purified 70S ribosomes by sucrose gradient centrifugation in a zonal rotor (Ti-15, Beckman, CA, USA) under dissociating conditions using buffer TAKM_3.5_ (50 mM Tris (pH 7.5), 70 mM NH_4_Cl, 30 mM KCl and 3.5 mM MgCl_2_), essentially as described^21^. Briefly, fractions containing 30S or 50S subunits were collected and pelleted in a Ti50.2 rotor at 50000 rpm over 12 hours. The resulting 30S pellets were resuspended in buffer TAKM7 (as TAKM_3.5_ but containing 7 mM MgCl_2_). The concentration of 30S subunits was determined by measuring the absorbance at 260 nm using an extinction coefficient of 63 pmol/AU_260 nm_.

#### Initiation factors

Cells harbouring pET 21 *(Kan* resistant) expression plasmids with cloned either *infA, infB* or *infC* (coding for IF1, 2 and 3, respectively) were grown in 6 L of LB medium supplemented with 30 μg/ml of kanamycin at 37°C until they reached an optical density of 0.8 OD_600_. A final concentration of 1 mM of IPTG was added to induce protein expression, leaving cells grow for 3 hours at same growth conditions. Cells were collected by centrifugation at 6000 RCF and resuspended in buffer A (50 mM Tris (pH 7.1), and 5% v/v glycerol) with 200 mM KCl. Prior to cell lysis 5 mM 2-mercaptoethanol, a protease tablet cocktail inhibitor (11836153001, Roche), 0,5 mM Pefabloc (11429868001, Roche) 1 mg/ml of lysozyme and few crystals of DNAse (DN25, Sigma-Aldrich) were added to the ice-cold suspension of unfrozen cells. Lysates were obtained using a Misonix 3000 sonicator (EW-04711-81, Misonix Inc) for 5 minutes with 25% amplitude for 10 s sonication followed by 20 s of pausing to avoid overheating. The lysate was centrifuged in a JA30.5 rotor for 30 min at 25000 rpm to remove cell debris. To dissociate initiation factors from the ribosomes, the concentration of KCl of the supernatant was increased to 0.7 M subsequently, ribosomes were sedimented by centrifugation in a Ti 50.2 rotor for 2 h at 50000 rpm, enriching the initiation factors in the supernatant. Supernatants containing either IF1 or IF3 were diluted with buffer A to reach a final concentration of 0.1M KCl. A HiTrap SP HP column (5 ml) (17-1151-01, GE Healthcare) was equilibrated with buffer A containing 0.1 M KCl prior to loading the clarified lysates. IF1 was eluted with a linear gradient of 0.05 – to 1 M KCl in buffer A. Fractions containing IF1 were identified using 18% acrylamide SDS-PAGE, pooled and concentrated using an Amicon centrifugation membrane with a 3 KDa cut-off (UFC900308, Merck). Size exclusion using a HiLoad 26/60 Superdex 75 prep grade column (17-1070-01, GE Healthcare) was necessary to further purify IF1 from contaminants of higher molecular weight. Typically, 0.5 ml of IF1 was loaded, separated with a flow rate of 1 ml/min and elution was monitored by absorbance at 290 nm. IF3 was purified similarly to IF1 using a cation exchange chromatography on a HiPrep CM FF 16/10 column (28-9365-42, GE Healthcare), however using a stronger gradient in Buffer A (0.1 to 1 M NH4Cl). A second chromatography step was unnecessarily, IF3 elutes with very high purity as observed using 15% SDS-PAGE.

Cell lysates containing IF2 were processed essentially as described for IF1 with the following modifications. Affinity chromatography on Hi-Trap His Column (54835, GE Healthcare) was used as a first step as a 6x His-tag was added at the amino terminal domain of IF2. A 50 to 300 mM imidazole gradient in Buffer A containing 300 mM KCl was used to elute IF2. Fractions containing the factor were pooled together and dialyzed overnight to buffer A containing 50 mM KCl prior to cation exchange chromatography. A 50 – 500 mM KCl gradient was used to elute IF2 from a 5 ml Hi-Trap SP HP column (17115201, GE Healthcare). Fractions were analysed by 8% acrylamide SDS-PAGE and those containing IF2 were pooled together. All three IFs were finally dialyzed in Storage Buffer (50 mM Tris (pH 7.1), 200 mM NH4Cl, 5 mM 2-mercaptoethanol and 10 % v/v Glycerol), small aliquots were flash frozen in liquid nitrogen and stored at −80°C.

#### Bpy-Met-tRNA^fMet^ (Bpy-tRNAi)

Met-tRNA^fMet^ was prepared essentially as described^30^. NHS ester BODIPY FL SSE dye (D6140, Invitrogen, or analogous) was used to label Met-tRNA^fMet^ at the amino group of the amino acid as follows. Met-tRNA^fMet^ was incubated in 50 mM Hepes-KOH (pH 8.5) with 3 mM of the dye in the dark. The reaction was stopped by adding 0.3 M potassium acetate (pH 5.0). Bpy-Met-tRNA^fMet^ was purified by three sequential ethanol precipitations and HPLC chromatography on a reverse phase C18 column with a 5 %-40 % ethanol gradient. The efficiency of labelling was determined by the molar stoichiometry of the compound measuring both, tRNA and dye absorbance.

#### mRNAs

DNA templates for mRNAs were amplified by PCR using the Maxima Hot Start Green PCR Master Mix (K1062, ThermoScientific) and corresponding primers (Table S1). Essentially, the reaction contained 20 ng of DNA template, 1 μM of each primer, Master Mix (containing Buffer, NTPs and polymerase) and deionized water. Primers were synthetized by Macrogen (South Korea). PCR products were purified using the GeneJET PCR Purification Kit (K0702, ThermoScientific) (Table S2). mRNAs were produced by *in vitro* transcription for 3 hours at 37°C. The reaction contained Transcription Buffer (40 mM Tris-HCl (pH 7.5), 15 mM MgCl_2_, 2 mM spermidine and 10 mM NaCl), 10 mM DL-Dithiothreitol (DTT), 2.5 mM NTPs, 5 mM guanosine monophosphate, 0.01u/μl inorganic pyrophosphatase, 2 U/μl T7 polymerase, 5 ng/μl DNA template and deionized water. Transcripts were then purified using Direct-ZolTM RNA MiniPrep (R2052, Zymo Research) and visualized by 8 M urea PAGE electrophoresis followed by staining in Methylene blue (Table S3).

#### Rel_Seq_

For preparation of ppGpp, we first purified the N-terminal fragment containing the 385 first amino acids of native RelSeq protein. The enzyme was purified from BL21 (DE3) cells transformed with pET21 plasmid encoding C-terminal 6×His-tagged fragment. Cells were grown in LB medium with 100 μg/ml ampicillin at 37 °C to an 0.6 OD_600_ and protein expression was induced with 1 mM IPTG with additional incubation for 3 h. Cells were collected by low speed centrifugation (6000 rpm × 20 min, JLA8.1 rotor). Cell pellets were resuspended in lysis buffer (20 mM Tris-HCl pH 7.9, 300 mM KCl, 5 mM MgCl_2_, 20% glycerol, 10 mM imidazole, 1 mM DTT, 280 μg/ml lysozyme, 0.1 mg/ml DNase I (DN25, Sigma-Aldrich) and 1 tablet of protease inhibitor cocktail (11836153001, Roche). Cells were opened using the EmulsiFlex-C3 (Avestin) and the cell debris was removed by the centrifugation (45000 rpm × 30 min, rotor Ti50.2). The supernatant was applied to a HisTrap FF (5 ml) column (17531901, GE Healthcare) for affinity chromatography. The column was washed with buffer HT (20 mM Tris-HCl pH 7.9, 5 mM MgCl_2_, 20% glycerol, 1 mM DTT) with 300 mM KCl and 10 mM imidazole and the protein was eluted by a linear gradient from 10 mM to 200 mM imidazole in the same buffer. The eluted product was dialyzed twice against the buffer HT and 450 mM KCl. The protein was concentrated by Amicon Ultra-15 Centrifugal Filters (10 KDa) (UFC901008, Merck) and diluted in buffer HT with 50 mM KCl for decreasing of KCl concentration to 90 mM. An anion-exchange chromatography step was used to further purify RelSeq. HiTrapQ (5 ml) column (17115301, GE Healthcare) was equilibrated with the buffer (20 mM Tris-HCl pH 9.5, 50 mM KCl, 5 mM MgCl_2_, 20% glycerol, 1 mM DTT) and the protein was eluted by a linear gradient from 50 mM to 750 mM KCl. Fractions containing RelSeq were analysed by 12% SDS-PAGE gel-electrophoresis, pooled, aliquoted and frozen in a liquid nitrogen. The enzyme activity of RelSeq was above 80%, as measured by ppGpp production over the sum of ppGpp plus GDP. *ppGpp*: preparative synthesis of the tetraphosphate was performed in buffer B (30 mM Tris-HCl pH 8.0, 100 mM NaCl, 10 mM MgCl_2_) using 10 mM ATP, 4 mM GDP and 50 μM RelSeq. The reaction was incubated for 40 min at 37 °C and stopped by phenol extraction. The water phase was loaded to a MonoQ 5/50 GL (1 ml) column (GE17-5166-01, GE Healthcare). ppGpp was eluted with a linear gradient from 0.5 mM to 600 mM LiCl in buffer C (25 mM Tris-HCl pH 8.3, 0. 5 mM EDTA). Fractions containing ppGpp were pooled and precipitated with 1.5 M LiCl and 2 volumes of absolute ethanol. The precipitate was harvested by centrifugation (16100 RFC × 20 min) and dissolved in water. ppGpp concentration was measured by UV absorbance at 253 nm using an extinction coefficient of 13700. Samples were aliquoted and stored at −20°C.

### Experimental conditions and analysis

30S complexes were reconstituted using pure subunits, IFs, tRNAi, mRNA and nucleotides. 30S subunits were reactivated by incubation in TAKM20 buffer (as TAKM3.5 but with 20 mM MgCl_2_) for 1 h at 37 °C prior to use. Generally, 30S ICs were formed by incubation at 37°C for 30 min in TAKM7 using 1 μM 30S subunits, 4 μM mRNA, 2 μM IF1, 1 μM IF2, 1.5 μM IF3, 0.5 μM Bpy-Met-tRNA^fMet^ and 0.5 mM guanosine nucleotides, unless otherwise stated in the results section or figure legends. Ligand titrations were prepared using serial dilutions by one to one mixing of complexes containing the highest concentration of the titrant with complexes lacking only the ligand under investigation. The number of reactions were setup to cover a wide range of concentrations.

MST was measured on 10 μl reactions using standard capillaries (MO-K022, NanoTemper Technologies) on a Monolith NT.115 (NanoTemper Technologies) at monochromatic LED with power input of 30% and IR laser power of 40%. The local temperature perturbation was expected to be less than 3 °C. All measurements were performed at room temperature (22 ± 2 °C). 3-5 replicates were measured for each investigated reaction to calculate mean and standard deviation. Day to day reproducibility was very high which allowed to report absolute thermophoresis values rather than to use normalized MST traces. *K_D_* were calculated using a hyperbolic or quadratic function using Graphpad Prism 6.0 (GraphPad Inc, USA) or software provided by NanoTemper. Same-site inhibitory constants were calculated using the *K_D_* values obtained for GTP and concentrations used by non-linear fitting using the one site competition function using Graphpad Prism 6.0 (GraphPad Inc, USA).

To measure 70S pre-IC formation, a SX-20 stopped flow instrument (Applied Photophysics, UK) was set up for light scattering recording with an excitation wavelength of 430 nm. Scattered light was measured at an angle of 90° without a filter. Pseudo first order conditions were approximated by using ≥3-fold excess of 50S subunits over 30S ICs. 0.2 μM 30S ICs were rapidly mixed in the instrument with 0.6 μM 50S at 25 °C. Bpy-tRNAi accommodation was measured essentially as described by^12^. Briefly, 0.2 μM 30S ICs were formed using Bpy-tRNAi and rapidly mixed with 0.6 μM 50S at 25 °C. Fluorescence was excited at 470 nm and measured after a 495 nm long pass filter. Individual traces (6-11 traces) were averaged and the resulting kinetic curve was approximated by single (F = A_0_ + A_1_*exp (-k^1^_app_*t)) or double (F = A_0_ + A_1_*exp (-k^1^_app_*t) + A_2_*exp (-k^2^_app_*t)) exponential fit using Graphpad Prism 6.0 software (GraphPad Inc, USA).

In order to visualize overall protein synthesis, we used a coupled transcription-translation cell-free system. Reactions were carried out in 10 μl of the purified components from the PURExpress *In Vitro* Protein Synthesis Kit (E6800S/L, NEB) with the corresponding templates. These reactions were incubated at 37°C for 2 hours. To asses ppGpp inhibitory capacity in translation, 0.15 μM of DNA coding for *mTuf*A or *mlnf*A were introduced in the described system in the absence or presence of 4, 2, 1, 0.5 or 0.25 mM ppGpp. After incubation, sample processing and fluorescent labelling were carried out as described by the Lumio™ Green Detection Kit (LC6090, ThermoScientific) for *in vitro* reactions. Finally, 20 μl of labelled samples were loaded into 20% SDS-PAGE gels and visualized under a Blue-light LED transilluminator with orange filter (Cleaver Scientific LTD). Gel quantification was performed by pixel densitometry analysis using ImageJ software^26^.

## Supplementary Information

Supplementary Text, experimental approach

Supplementary Figures 1 to 8

Supplementary Tables 1 to 3

## Acknowledgments

We are thankful to Vasili Hauryliuk for the construct coding for RelSeq and to Claudio Gualerzi for sharing plasmids coding for model mRNAs. Experiments on dynamics of the ribosomal complexes were supported by Russian Science Foundation Grant 17-14-01416 (to ALK), experiments on various mRNAs were supported by Russian Foundation for Basic Research grant 17-00-00368 (to ALK), by InnóvatePerú grants 382-PNICP-PIBA-2014 and 297-INNOVATEPERU-EC-2016 (to PM), and by the Fondecyt grant 154-2017-Fondecyt (to PM). The Monolith NT.115 equipment used for MST measurements was provided by Nanotemper Technologies RUS LLC.

## Author Contributions

ALK and PM conceived the project; DSV, VZ, PK and EM performed the experiments; DSV, AP, ALK, and PM analysed the data; AP, ALK and PM wrote the manuscript with the input of all authors.

