## Supplementary Information for "How the initiating ribosome copes with (p)ppGpp to translate mRNAs"

This PDF file includes:

- Supplementary Text, experimental approach
- Supplementary figures 1 to 8
- Supplementary tables 1 to 3

### Experimental approach

To monitor translation initiation as a function of guanosine nucleotides, mRNAs and IFs, we modified tRNAi at the amino acyl moiety with the fluorescent dye BODIPY-FL (30) (Bpy-tRNAi) (Supplementary Fig. 1). The equilibrium binding of Bpy-tRNAi to 30S ICs was studied by Microscale Thermophoresis (MST). MST allows to assess molecular drifts as a function of equilibrium perturbations by small changes in temperature (Supplementary Fig. 1). Upon equilibrium perturbations, the observed labelled compound migration patterns are related to bound and unbound states of the reactants, allowing to determine dissociation constants of the interaction (Supplementary Fig. 1). By observing the bound state of Bpy-tRNAi to the 30S subunit, MST could be measuring either complex, the 30S pre-IC and 30S IC as they are identical in terms of ligand composition. However, decoding of the start codon in the 30S IC was shown to increase the kinetic stability of tRNAi (19). Titrations with an mRNA lacking a start codon (30S pre-IC) showed negligible thermophoresis signal (Supplementary Fig. 2), indicating that MST monitors the formation of the 30S IC rather than the 30S pre-IC. To further validate this observation, we used a Bpy-Phe-tRNA$^{Phe}$ on 30S pre-ICs and observed an increase of thermophoresis only for complexes programmed with the corresponding UUC codon at the start site (Supplementary Fig. 2).

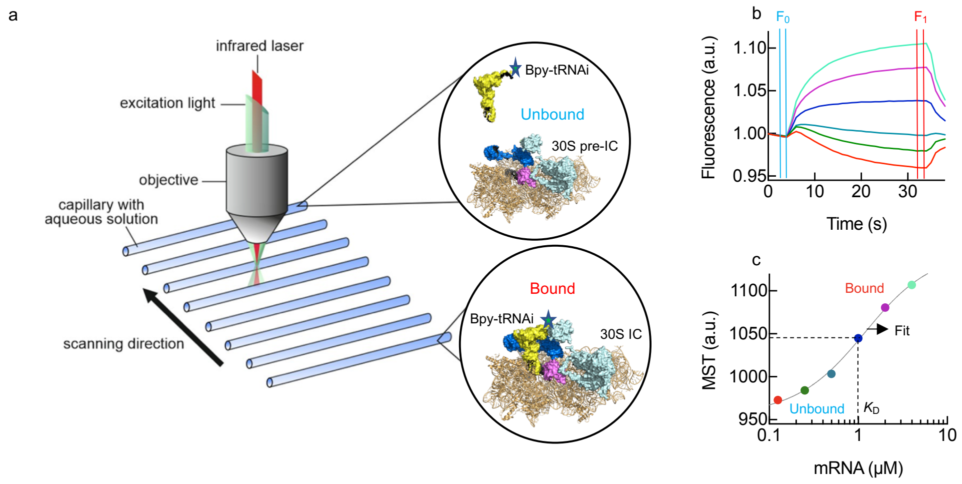

**Supplementary Figure 1**

Experimental approach to determine the binding of Bpy-tRNAi to 30S IC using Microscale Thermophoresis. **(a)** Instrument concept. **(b)** Fluorescence measurements on time as function of ligand concentration. Blue vertical lines indicate the fluorescence interval prior to the equilibrium perturbation (Cold) while vertical red lines indicate the fluorescence interval upon reaching the new equilibrium (Hot). Colored traces represent different concentrations of the ligands. The ratio between fluorescence hot and cold fluorescence (times 1000) is related to the thermophoresis shift (MST). **(c)** The dependency of MST from ligand concentration allows to estimate $K_D$ constants of the interaction using a quadratic function and non-linear regression fitting. MST measurements (full circles) are indicated in the same colors as their respective time traces (b).

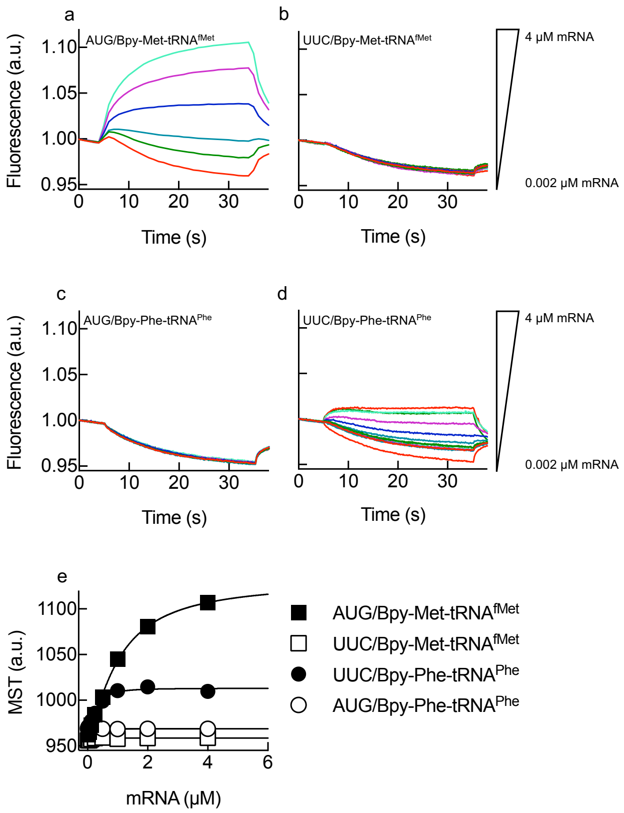


**Supplementary Figure 2**

Step assignment for 30S IC formation by MST measurements. Time dependencies of fluorescence measurements for Bpy-tRNAi interacting with 30S ribosomal complexes programmed with mMF1 containing AUG **(a)** or UUC **(b)** initiation codon. Time traces of Bpy-Phe-tRNA$^{Phe}$ binding to 30S complexes using mMF1 containing AUG **(c)** or UUC **(d)** as initiation codons. Colors represent increasing concentrations of mRNA (from 2 nM to 4 µM). **(e)** Comparison of all four combinations of start codons and labeled tRNAs for MST dependencies on mRNA concentration. Continuous lines indicate non-linear regression fittings. 3-5 measurements were performed, mean and standard deviation are plotted.

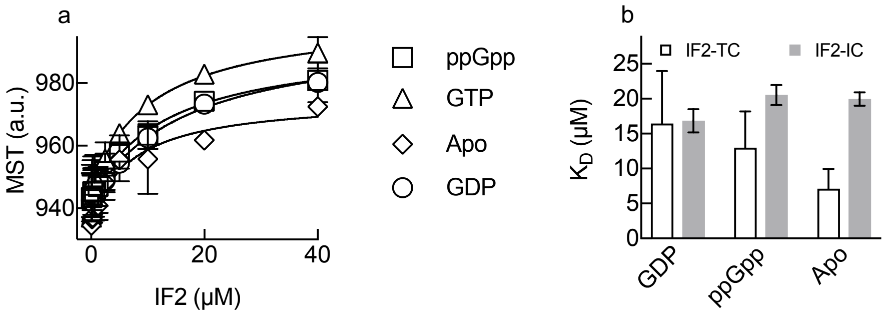


**Supplementary Figure 3**

IF2 to Bpy-tRNAi ternary complex formation by MST measurements. **(a)** IF2 concentrations dependencies for Bpy-tRNAi binding in the presences of 0.5 mM of the indicated guanosine nucleotides. **(b)** Comparison of the calculated $K_D$ from interactions measured in (a) with a similar IF2 titration performed in the complete 30S IC complex (Fig. 2a). 3-5 measurements were performed, mean and standard deviation are plotted.

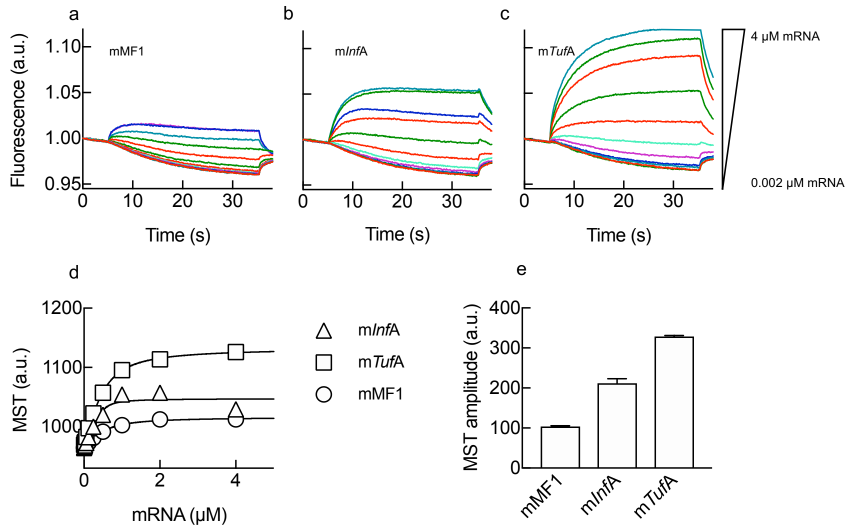


**Supplementary Figure 4**

Formation of 30S IC as a function of the mRNA, model or natural. **(a)** Time courses of 30S IC formation for increasing concentrations of the model mMF1. **(b)** Time courses of mInfA or **(c)** mTufA dependent 30S IC formation. Colors represent increasing concentrations of mRNA (from 2 nM to 4 μM). **(d)** MST dependence on the mRNA concentrations for all three tested templates. Continuous lines show the non-linear fitting. 3-5 measurements were performed, mean and standard deviation are plotted. **(e)** Comparison of MST amplitudes as an indicator of the efficiency of 30S IC formation with all three mRNAs.

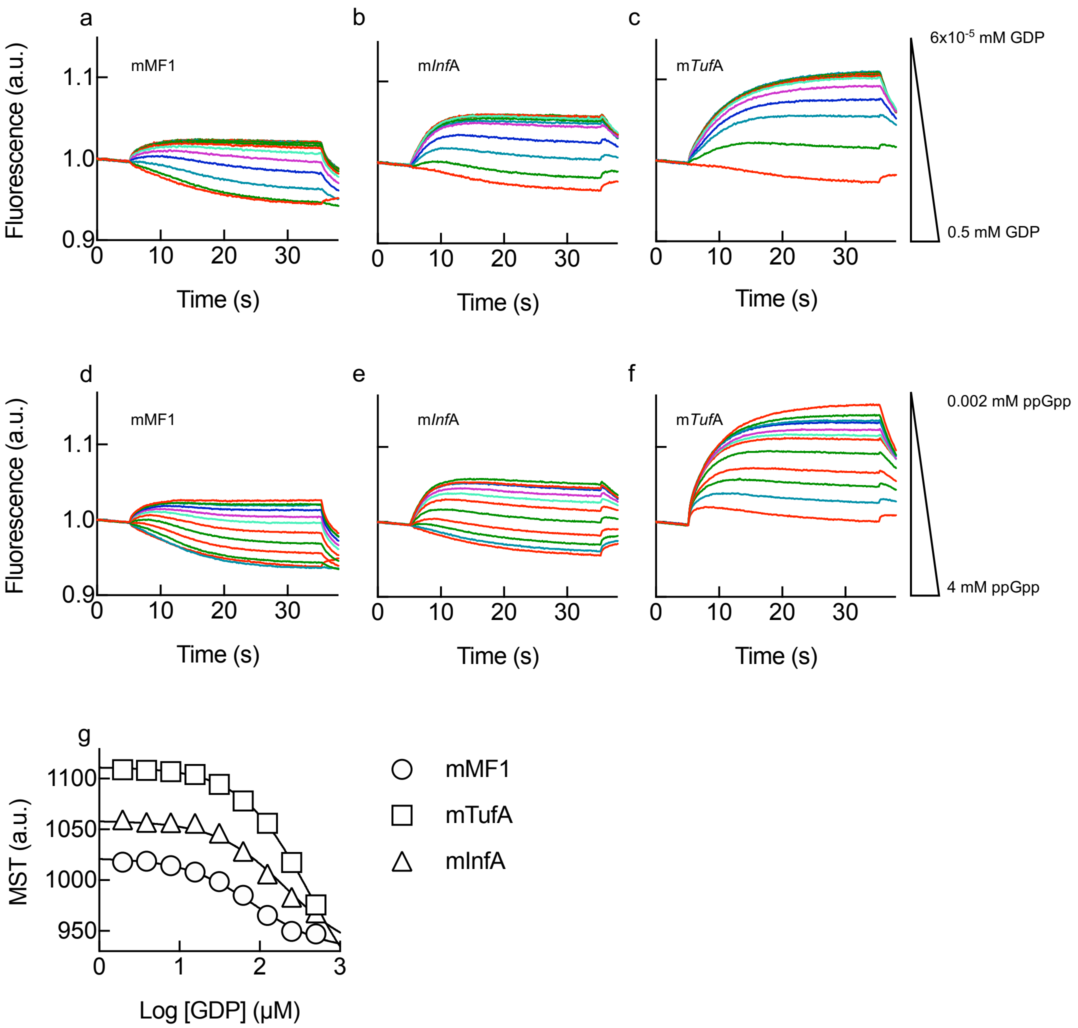


**Supplementary Figure 5**

Inhibition of 30S IC formation by GDP or ppGpp competition with GTP (Related to Fig. 3). **(a)**, **(b)** and **(c)** show time courses of thermophoresis for mMF1, mInfA and mTufA, respectively, at increasing concentrations of GDP competing with 50 μM GTP used for 30S IC formation. **(d)**, **(e)** and **(f)** show time courses of thermophoresis for mMF1, mInfA and mTufA, respectively, at increasing concentrations of ppGpp competing with 50 μM GTP used for 30S IC formation. **(g)** MST dependency as a function of the logarithmic of GDP concentration. Symbols are as indicated, 3-5 measurements were performed, mean and standard deviation are plotted. Continuous lines show non-linear regression fittings with a same-site inhibition model.

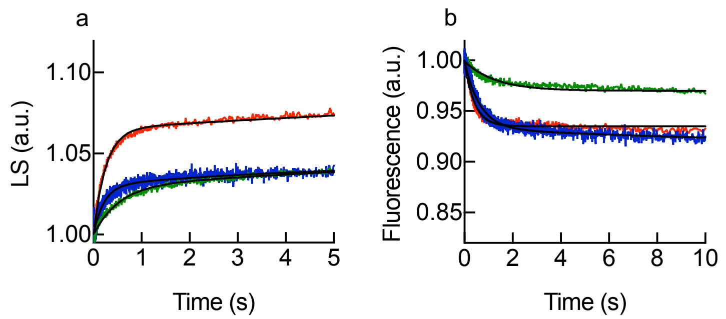


**Supplementary Figure 6**

Inhibitory effect of 70S pre-IC and 70S IC formation by GDP competing with GTP for IF2 (related to Fig. 3). 30S IC containing 20 µM GTP and 200 µM GDP, programmed with mMF1 (blue), mInfA (green) or mTufA (red), were rapidly mixed with 50S subunits in a stopped-flow apparatus and scattered light **(a)** or Bpy-tRNAi fluorescence **(b)** were monitored over time as described in Fig. 4. 7-10 individual replicates were recorded. Non-linear regression fitting with exponential functions is shown as continuous black lines.

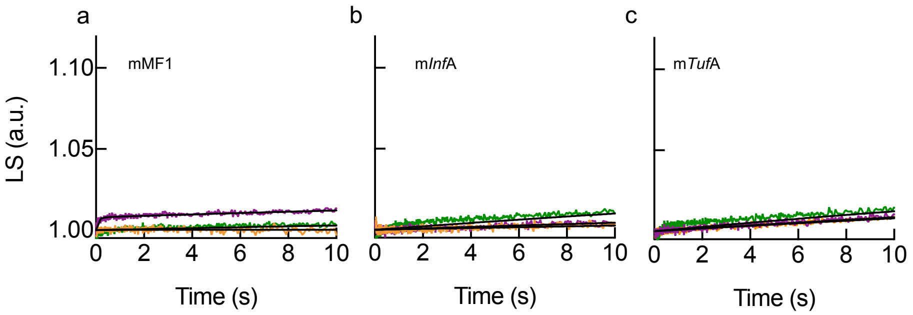


**Supplementary Figure 7**

Guanosine nucleotide dependencies for 70S pre-IC formation under noncompetitive conditions (related to Fig. 4). Time courses of 70S pre-ICs formation in the absence of any nucleotide (purple) or in the presence of 0.2 mM of either GDP (green) or ppGpp (orange). 30S ICs were programmed with either mMF1 **(a)**, mInfA **(b)** or mTufA **(c)**. 70S pre-IC was measured by the scattered light with a stopped-flow apparatus. 7-10 individual replicates were recorded. Non-linear regression fitting is shown as continuous black lines.

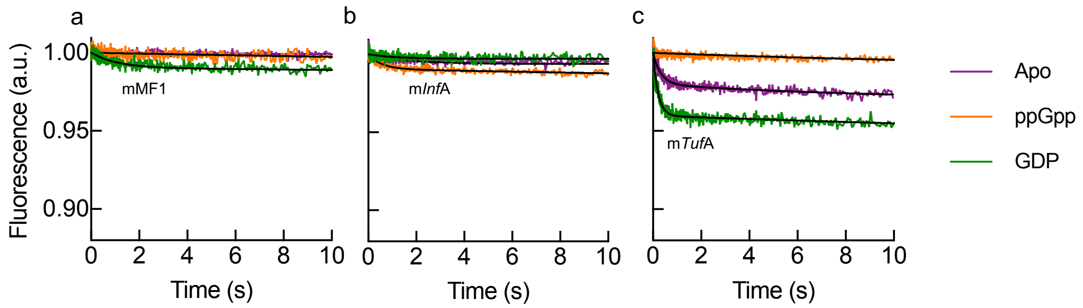


**Supplementary Figure 8**

Guanosine nucleotide dependencies for 70S IC formation as monitored by Bpy-tRNAi accommodation under noncompetitive conditions (related to Fig. 4). Time courses of 70S ICs formation in the absence of any nucleotide (purple) or in the presence of 0.2 mM of either GDP (green) or ppGpp (orange). 30S ICs were programmed with either mMF1 **(a)**, mInfA **(b)** or mTufA **(c)**. 70S IC was measured by fluorescence change of Bpy-tRNAi with a stopped-flow apparatus. 7-10 individual replicates were recorded. Non-linear regression fitting is shown as continuous black lines.

**Supplementary Table 1**

Primer sequences for DNA template amplification. TufA and InfA are naturally encoded in the E. coli genome while MF1 coding sequence is cloned in a pTZ18R plasmid. Primers for MF1 correspond to the T7 promoter and terminator sequences. The sequence coding for a tetra cysteine motif, added as an overhang sequence to the reverse primer, is indicated with the L suffix. A T7 promoter overhang was added to the forward primer to allow in vitro transcription.

| Name | Forward primer (5'→3') | Reverse primer (5'→3') | DNA (bp) | mRNA (nt) |
|---|---|---|---|---|
| TufA | TAATACGACTCACTATAGGTTCTATCGCCTTTAAAGAAGGC | GCATGTTAGGTGATTGCAGCGGTCAGAG | 514 | 497 |
| TufA_L | TAATACGACTCACTATAGGTTCTATCGCCTTTAAAGAAGGC | GCATGTTAGCAGCAGCCCGGGCAGCAGGTGATTGCAGCGGTCAGAG | 540 | 523 |
| InfA | TAATACGACTCACTATAGGCGCAGAGTTGGTTACGCTC | GCATGTCACAGTAACCACGTGACCG | 361 | 344 |
| InfA_L | TAATACGACTCACTATAGGCGCAGAGTTGGTTACGCTC | GCATGTCAGCAGCAGCCCGGGCAGCACAGTAACCACGTGACCG | 387 | 370 |
| MF1 | CGAATTTAATACGACTCACTATAGG | GCTTGCATGCCTGCAGACGCA | 121 | 93 |

**Supplementary Table 2**

Gene sequences used for mRNA synthesis. Both, TufA and InfA contain two promoters. P1 was used in this study (red) for both natural mRNAs while mMF1 uses a T7 promoter encoded in the plasmid (red). Blue indicates 5' untranslated regions (UTR). Brown indicates the coding sequences corresponding to final DNA templates used for in vitro transcription. Lower case indicates the start codon. Underlined sequences indicate priming regions for PCR amplifications (Table S1). PCR of the template p022 generates a short coding region for in vitro transcription of the mMF1.

| Name | DNA sequences |
|---|---|
| TufA | TTCTATCGCCTTTAAAGAAGGCTTTAAGAAAGCGAAACCAGTTCTGCTTGAGCCGATCAT<br>GAAGGTTGAAGTAGAAACTCCGGAAGAGAACACCGGTGACGTTATCGGTGACTTGAGCC<br>GTCGTCGTGGTATGCTCAAAGGTCAGGAATCTGAAGTTACTGGCGTTAAGATCCACGCTG<br>AAGTACCGCTGTCTGAAATGTTCGGATACGCAACTCAGCTGCGTTCTCTGACCAAAGGTC<br>GTGCATCATACACTATGGAATTCCTGAAGTATGATGAAGCGCCGAGTAACGTTGCTCAGG<br>CCGTAATTGAAGCCCGTGGTAAATAAGCCTAAGGGTTAATACCAAAGTCCCGTGCTCTCT<br>CCTGAAGGGGAGAGCACTATAGTAAGGAATATAGCCgtgTCTAAAGAAAAATTTGAACGT<br>ACAAAACCGCACGTTAACGTTGGTACTATCGGCCACGTTGACCACGGTAAAACTACTCTG<br>ACCGCTGCAATCACCACCGTACTGGCTAAAACCTACGGCGGTGCTGCTCGTGCATTCGAC<br>CAGATCGATAACGCGCCGGAAGAAAAAGCTCGTGGTATCACCATCAACACTTCTCACGTT<br>GAATACGACACCCCGACCCGTCACTACGCACACGTAGACTGCCCGGGGCACGCCGACTAT<br>GTTAAAAACATGATCACCGGTGCTGCTCAGATGGACGGCGCGATCCTGGTAGTTGCTGCG<br>ACTGACGGCCCGATGCCGCAGACTCGTGAGCACATCCTGCTGGGTCGTCAGGTAGGCGTT<br>CCGTACATCATCGTGTTCCTGAACAAATGCGACATGGTTGATGACGAAGAGCTGCTGGAA<br>CTGGTTGAAATGGAAGTTCGTGAACTTCTGTCTCAGTACGACTTCCCGGGCGACGACACT<br>CCGATCGTTCGTGGTTCTGCTCTGAAAGCGCTGGAAGGCGACGCAGAGTGGGAAGCGAA<br>AATCCTGGAACTGGCTGGCTTCCTGGATTCTTATATTCCGGAACCAGAGCGTGCGATTGA<br>CAAGCCGTTCCTGCTGCCGATCGAAGACGTATTCTCCATCTCCGGTCGTGGTACCGTTGTT<br>ACCGGTCGTGTAGAACGCGGTATCATCAAAGTTGGTGAAGAAGTTGAAATCGTTGGTATC<br>AAAGAGACTCAGAAGTCTACCTGTACTGGCGTTGAAATGTTCCGCAAACTGCTGGACGAA<br>GGCCGTGCTGGTGAGAACGTAGGTGTTCTGCTGCGTGGTATCAAACGTGAAGAAATCGAA<br>CGTGGTCAGGTACTGGCTAAGCCGGGCACCATCAAGCCGCACACCAAGTTCGAATCTGAA<br>GTGTACATTCTGTCCAAAGATGAAGGCGGCCGTCATACTCCGTTCTTCAAAGGCTACCGT<br>CCGCAGTTCTACTTCCGTACTACTGACGTGACTGGTACCATCGAACTGCCGGAAGGCGTA<br>GAGATGGTAATGCCGGGCGACAACATCAAAATGGTTGTTACCCTGATCCACCCGATCGCG<br>ATGGACGACGGTCTGCGTTTCGCAATCCGTGAAGGCGGCCGTACCGTTGGCGCGGGCGTT<br>GTTGCTAAAGTTCTGGGCTAA |
| InfA | CGCAGAGTTGGTTACGCTCATTACCCCGCTGCCGATAAGGAATTTTTCGCGTCAGGTAAC<br>GCCCATCGTTTATCTCACCGCTCCCTTATACGTTGCGCTTTTGGTGCGGCTTAGCCGTGTGT |

| | |
|---|---|
| | TTTCGGAGTAATGTGCCGAACCTGTTTGTTGCGATTTAGCGCGCAAATCTTTACTTATTTA<br>CAGAACTTCGGCATTATCTTGCCGGTTCAAATTACGGTAGTGATACCCCAGAGGATTAGat<br>gGCCAAAGAAGACAATATTGAAATGCAAGGTACCGTTCTTGAAACGTTGCCTAATACCAT<br>GTTCCGCGTAGAGTTAGAAACGGTCACGTGGTTACTGACACACATCTCCGGTAAAATGCG<br>CAAAAACTACATCCGCATCCTGACGGGCGACAAAGTGACTGTTGAACTGACCCCGTACGA<br>CCTGAGCAAAGGCCGCATTGTCTTCCGTAGTCGCTGA |
| p022<br>(MF1) | CGAATTTTAATACGACTCACTATAGGGAATTCAAAAATTTAAAAGTTAACAGGTATACAT<br>ACT(atg/ttc)TTTACGATTACTACGATCTTCTTCACTTAATGCGTCTGCAGGCATGCAAGCTA<br>AAAAAAAAAAAAAAAAAAAAAAAAAAGCTTGGCACTGGCCGTCGTTTTACAACGTCGTG<br>ACTGGGAAAACCCTGGCGTTACCCAACTTAATCGCCTTGCAGCACATCCCCCTTTCGCCA<br>GCTGGCGTAATAGCGAAGAGGCCCGCACCGATCGCCCTTCCCAACAGTTGCGCAGCCTGA<br>ATGGCGAATGGAAATTGTAAGCGTTAATATTTTGTTAAAATTCGCGTTAAATTTTTGTTAA<br>ATCAGCTCATTTTTTAACCAATAGGCCGAAATCGGCAAAATCCCTTATAAATCAAAAGAA<br>TAGACCGAGATAGGGTTGAGTGTTGTTCCAGTTTGGAACAAGAGTCCACTATTAAAGAAC<br>GTGGACTCCAACGTCAAAGGGCGAAAAACCGTCTATCAGGGCGATGGCCCACTACGTGA<br>ACCATCACCCTAATCAAGTTTTTTGGGGTCGAGGTGCCGTAAAGCACTAAATCGGAACCC<br>TAAAGGGAGCCCCCGATTTAGAGCTTGACGGGGAAAGCCGGCGAACGTGGCGAGAAAGG<br>AAGGGAAGAAAGCGAAAGGAGCGGGCGCTAGGGCGCTGGCAAGTGTAGCGGTCACGCT<br>GCGCGTAACCACCACACCCGCCGCGCTTAATGCGCCGCTACAGGGCGCGTCAGTGGCACT<br>TTTCGGGGAAA |

**Supplementary Table 3**

mRNA sequences for MST and in vitro translation (L) analysis. Lower case indicates the start codon. For mMF1 parenthesis indicate alternative start codons.

| Name | mRNA Sequence |
|---|---|
| mTufA | GGUUCUAUCGCCUUUAAAGAAGGCUUUAAGAAAGCGAAACCAGUUCUGCUUGAG<br>CCGAUCAUGAAGGUUGAAGUAGAAACUCCGGAAGAGAACACCGGUGACGUUAUC<br>GGUGACUUGAGCCGUCGUCGUGGUAUGCUCAAAGGUCAGGAAUCUGAAGUUACU<br>GGCGUUAAGAUCCACGCUGAAGUACCGCUGUCUGAAAUGUUCGGAUACGCAACUC<br>AGCUGCGUUCUCUGACCAAAGGUCGUGCAUCAUACACUAUGGAAUUCCUGAAGUA<br>UGAUGAAGCGCCGAGUAACGUUGCUCAGGCCGUAAUUGAAGCCCGUGGUAAAUA<br>AGCCUAAGGGUUAAUACCAAAGUCCCGUGCUCUCUCCUGAAGGGGAGAGCACUAU<br>AGUAAGGAAUAUAGCCgugUCUAAAGAAAAAUUUGAACGUACAAAACCGCACGUU<br>AACGUUGGUACUAUCGGCCACGUUGACCACGGUAAAACUACUCUGACCGCUGCAA<br>UCACCA |
| mTufA_L | GGUUCUAUCGCCUUUAAAGAAGGCUUUAAGAAAGCGAAACCAGUUCUGCUUGAG<br>CCGAUCAUGAAGGUUGAAGUAGAAACUCCGGAAGAGAACACCGGUGACGUUAUC<br>GGUGACUUGAGCCGUCGUCGUGGUAUGCUCAAAGGUCAGGAAUCUGAAGUUACU<br>GGCGUUAAGAUCCACGCUGAAGUACCGCUGUCUGAAAUGUUCGGAUACGCAACUC<br>AGCUGCGUUCUCUGACCAAAGGUCGUGCAUCAUACACUAUGGAAUUCCUGAAGUA<br>UGAUGAAGCGCCGAGUAACGUUGCUCAGGCCGUAAUUGAAGCCCGUGGUAAAUA<br>AGCCUAAGGGUUAAUACCAAAGUCCCGUGCUCUCUCCUGAAGGGGAGAGCACUAU<br>AGUAAGGAAUAUAGCCgugUCUAAAGAAAAAUUUGAACGUACAAAACCGCACGUU<br>AACGUUGGUACUAUCGGCCACGUUGACCACGGUAAAACUACUCUGACCGCUGCAA<br>UCACCAUGCUGCCCGGGCUGCUGCUGACAUGC |
| mInfA | GGCGCAGAGUUGGUUACGCUCAUUACCCCGCUGCCGAUAAGGAAUUUUUCGCGUC<br>AGGUAACGCCCAUCGUUUAUCUCACCGCUCCCUUAUACGUUGCGCUUUUGGUGCG<br>GCUUAGCCGUGUGUUUUCGGAGUAAUGUGCCGAACCUGUUUGUUGCGAUUUAGC<br>GCGCAAAUCUUUACUUAUUUACAGAACUUCGGCAUUAUCUUGCCGGUUCAAAUUA<br>CGGUAGUGAUACCCCAGAGGAUUAGaugGCCAAAGAAGACAAUAUUGAAAUGCAA<br>GGUACCGUUCUUGAAACGUUGCCUAAUACCAUGUUCCGCGUAGAGUUAGAAAACG<br>GUCACGUGGUUACUG |
| mInfA_L | GGCGCAGAGUUGGUUACGCUCAUUACCCCGCUGCCGAUAAGGAAUUUUUCGCGUC<br>AGGUAACGCCCAUCGUUUAUCUCACCGCUCCCUUAUACGUUGCGCUUUUGGUGCG<br>GCUUAGCCGUGUGUUUUCGGAGUAAUGUGCCGAACCUGUUUGUUGCGAUUUAGC<br>GCGCAAAUCUUUACUUAUUUACAGAACUUCGGCAUUAUCUUGCCGGUUCAAAUUA<br>CGGUAGUGAUACCCCAGAGGAUUAGaugGCCAAAGAAGACAAUAUUGAAAUGCAA<br>GGUACCGUUCUUGAAACGUUGCCUAAUACCAUGUUCCGCGUAGAGUUAGAAAACG<br>GUCACGUGGUUACUGUGCUGCCCGGGCUGCUGCUAACAUGC |

| | |
|---|---|
| mMF1 | GGGAAUUCAAAAAUUUAAAAGUUAACAGGUAUACAUACU(aug/uuc)UUUACGAUUA<br>CUACGAUCUUCUUCACUUAAUGCGUCUGCAGGCAUGCAAGCGGGAAUUCAAAAAU<br>UUAAAAGUUAACAGGUAUACAUACU |
